# From pixels to pleasure: visual features explain dynamic aesthetic experiences across distinct movie content

**DOI:** 10.64898/2026.04.14.718404

**Authors:** Mustafa Alperen Ekinci, Nina Buhlmann, Daniel Kaiser

**Affiliations:** Neural Computation Group, Justus Liebig University Giessen, 35392 Giessen, Germany; Center for Mind, Brain and Behavior (CMBB), Universities of Giessen, Marburg, and Darmstadt, 35032 Marburg, Germany; Center for Applied Computer Science and Data Science (ZAD), Justus Liebig University Giessen, 35392 Giessen, Germany; Cluster of Excellence “The Adaptive Mind”, Universities of Giessen, Marburg, and Darmstadt, 35392 Giessen, Germany

**Keywords:** Empirical aesthetics, visual beauty, art appreciation, movie watching, deep neural network

## Abstract

Aesthetic experiences in everyday life unfold under continuously changing visual input. Although these experiences clearly depend on the observer and context, they are partly explained by the visual features of the input. Here, we investigated how well a combination of visual features predicts dynamic aesthetic experiences during naturalistic and artistic movie watching. In two experiments, participants continuously rated the aesthetic appeal of either the nature documentary *Home* or the animated art-style movie *Loving Vincent*. We modeled moment-to-moment ratings using image-computable visual features extracted from each movie frame, including visual fluency, color and motion statistics, and symmetry. Linear models trained on these features reliably predicted aesthetic ratings for new movie parts, both within and across observers, pointing to shared perceptual influences on aesthetic experiences. Model comparisons showed that visual fluency and color-related features were most informative for predicting aesthetic experience in both movies. Critically, models trained on one movie could reliably predict aesthetic appeal ratings in the other movie, despite the movies’ remarkably different content and styles. Color features were most informative for cross-movie prediction. We conclude that visual features shape dynamic and naturalistic aesthetic experiences, and that the mapping of visual features onto aesthetic appeal is stable across observers and different movie content.

## Introduction

Complex and dynamic visual stimuli are often associated with aesthetic experiences. In cinema, certain movie scenes captivate us with their beauty, such as the breathtaking New Zealand landscapes in *The Lord of the Rings* trilogy, whereas others remain visually unremarkable. This observation extends beyond cinema: hiking through those same New Zealand landscapes should be equally beautiful, while our everyday walk to work is often less so.

What factors drive these differences in aesthetic appeal? Research on visual aesthetics has linked aesthetic experiences to sensory and perceptual processing: some visual features are reliably connected to perceiving beauty, while others predispose reduced beauty (Brielmann & Dayan, 2022; Chatterjee, 2011; Fechner, 1876; Iigaya et al., 2021; Leder et al., 2004; Leder & Nadal, 2014; Palmer et al., 2013; Redies, 2015). For example, people tend to prefer curved over sharp-edged objects (Corradi et al., 2019), and symmetric over asymmetric faces (Rhodes et al., 1998). While we have learned a lot about which features are connected to the perception of beauty, the current literature remains highly fragmented on two grounds: prior work has often focused on (i) individual features in isolation (Enquist & Arak, 1994; Silvia & Barona, 2009) and (ii) just one type of visual content (Kirk et al., 2009; Leder & Nadal, 2014; Vessel & Rubin, 2010).

First, aesthetic preferences have largely been examined with respect to one candidate visual feature at a time (Mather et al., 2023). Such studies showed that preferences are influenced by low-level properties like luminance, contrast, and color (Graham et al., 2016; Schloss & Palmer, 2011; Swartz et al., 2024), as well as structural mid- and higher-level features like symmetry, complexity, or curvature (Huang et al., 2018; Van Geert & Wagemans, 2020; Vartanian et al., 2024). On a configural level, Gestalt principles (Van Geert & Wagemans, 2024) and visual fluency (Reber et al., 2004; Oppenheimer, 2008) have been connected to perceived beauty. Yet most of these studies only looked at one or relatively few of these features at a time, rendering it unclear how well aesthetic appeal can be modeled once all candidate features are considered.

Second, research on aesthetic preferences has typically examined the importance of visual features within a single type of visual content, such as artworks (Leder et al., 2006; Vessel & Rubin, 2010; Vessel et al., 2012), objects (Hekkert et al., 2003; Carbon, 2010), faces (Jones et al., 2007; Perrett et al., 1999), or natural scenes (Dhar et al., 2011; Farzanfar & Walther, 2023). This domain-specific focus has limited insight into how aesthetic appeal generalizes across different visual content, and whether the inferred importance of certain feature configurations holds true when different types of complex visual stimuli are considered.

As a consequence, we currently know little about how multiple visual properties jointly contribute to aesthetic judgments and how such feature-based accounts generalize across different content. Here, we address these limitations by examining a comprehensive set of visual features across two very different movies. In two experiments, our participants either watched the nature documentary *Home*, which includes a variety of natural scenes with varying levels of beauty (Experiment 1), or the animated art-style movie *Loving Vincent*, an animated depiction of Van Gogh’s artistic work and biography (Experiment 2). In both experiments, they continuously rated the movies’ aesthetic appeal on a moment-to-moment level (Fig. 1). By modeling continuous aesthetic ratings from a range of image-computable features extracted on a frame-to-frame level (Fig. 2), we test whether (i) visual features reliably predict aesthetic experiences in free-flowing movies, (ii) how these predictions generalize across observers, and (iii) how they generalize across the two different movies.

**Figure 1.**
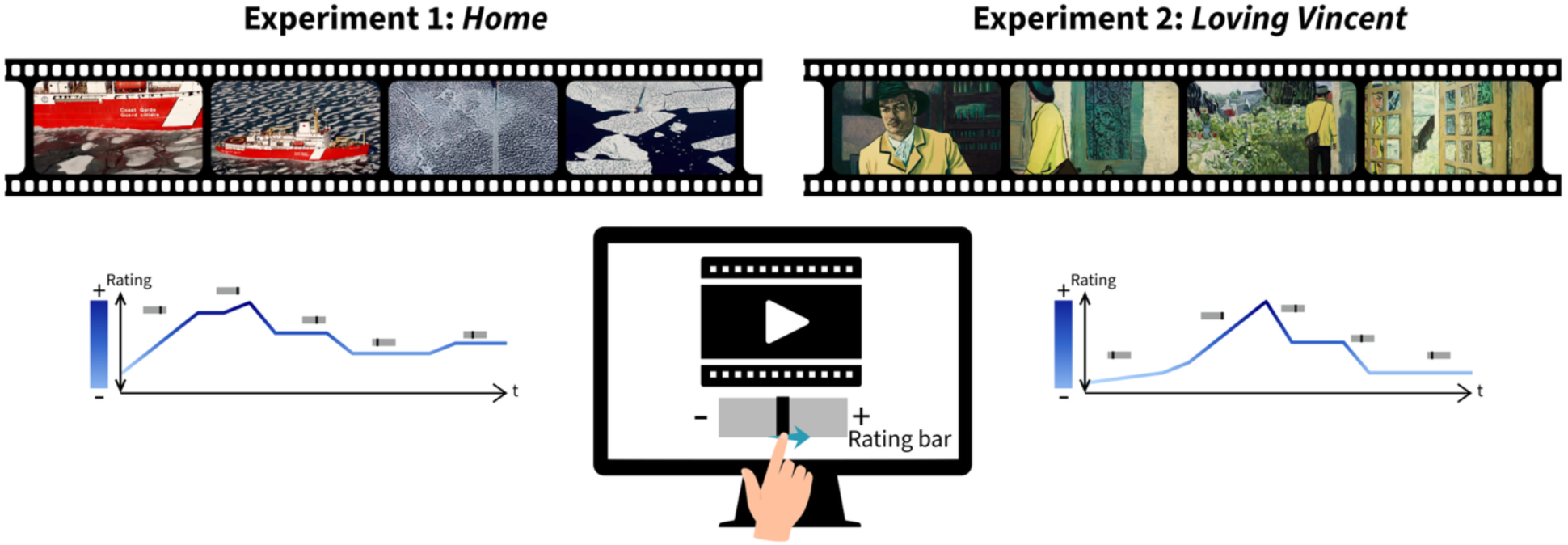
Paradigm. Participants continuously rated the aesthetic appeal of the movies *Home* (Experiment 1) or *Loving Vincent* (Experiment 2) using a slider bar.

**Figure 2.**
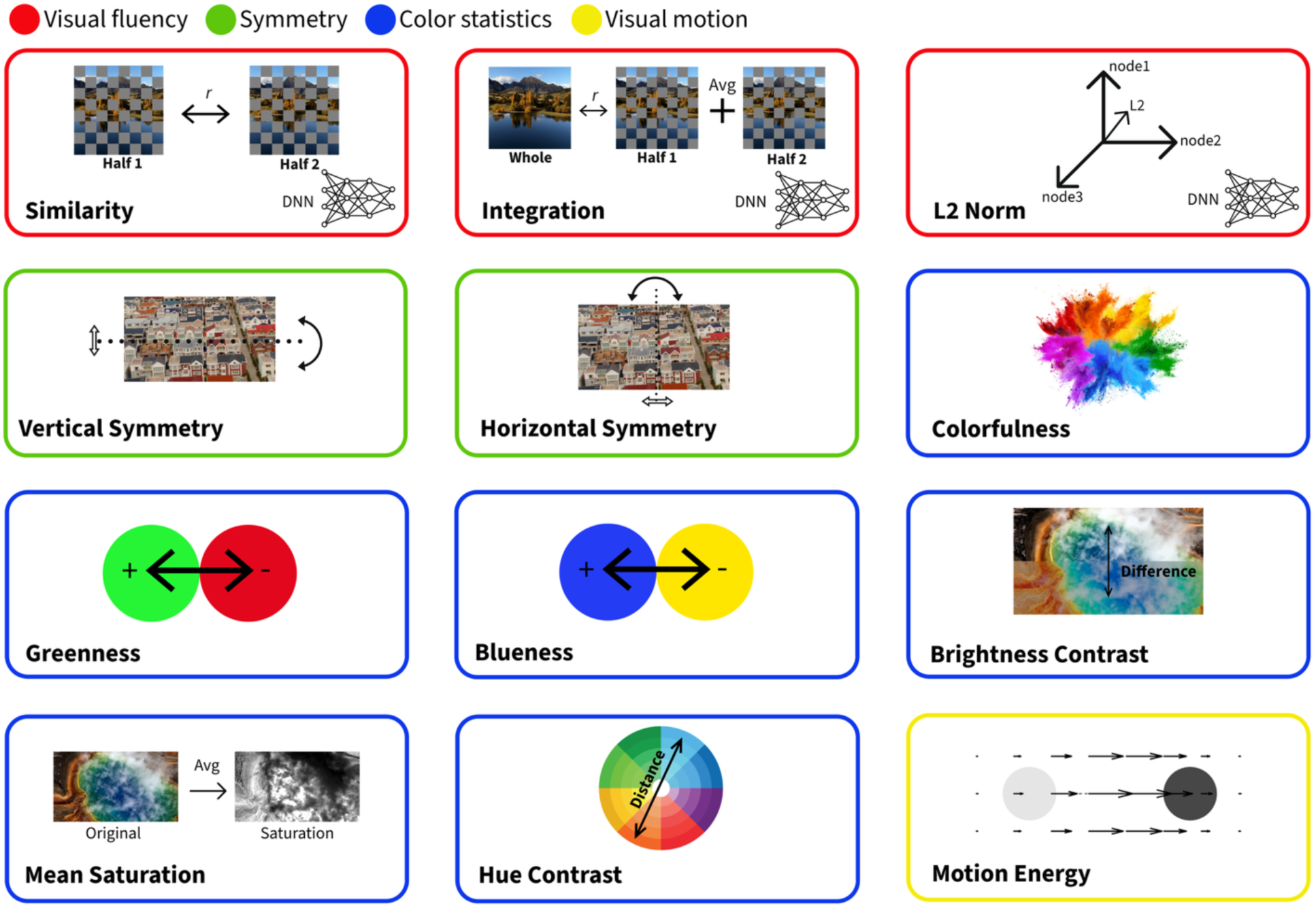
Image-computable predictors. We extracted twelve visual predictor metrics from individual movie frames. Predictors comprised four groups, covering aspects of visual fluency (similarity, integration, and L2 norm), symmetry (vertical and horizontal symmetry), color statistics (colorfulness, greenness, blueness, mean saturation, brightness contrast, and hue contrast), and visual motion (motion energy).

## Results

In Experiment 1, a total of 37 participants viewed the naturalistic movie *Home*, which was divided into nine segments (mean duration = 505.1 seconds). In Experiment 2, 30 participants viewed the artistic movie *Loving Vincent*, which was divided into eight segments (mean duration = 581.3 seconds).

For both movies, we extracted visual feature descriptors from each frame and resampled them to a one-second resolution to align with moment-to-moment aesthetic appeal ratings. These visual features were used as predictors to model aesthetic appeal ratings. A total of twelve predictors were extracted, split into four groups (Fig. 2): First, we computed visual fluency predictors (similarity, integration, and L2 norm) from activations in a deep neural network model (VGG16) by correlating representations of parts and wholes in different ways (Fig. S4). These predictors describe how much different regions of an image resemble one another (similarity), how well these regions combine into a coherent whole (integration), and the overall activation elicited by an image (L2 norm). Greater similarity and integration, as well as a lower L2 norm, have previously been associated with greater aesthetic appeal (Lin et al., 2025; Nara & Kaiser, 2024). Second, symmetry predictors (horizontal and vertical symmetry) were computed by correlating pixel values with their reflections across all possible symmetry axes (Damiano et al., 2023). Third, color predictors (colorfulness, greenness, blueness, mean saturation, hue contrast, and brightness contrast) were computed from pixel-wise color distributions (Hasler & Suesstrunk, 2003; Iigaya et al., 2021). These color features capture how much colors vary across the image, which colors are most prominent, and how much contrast exists both between different colors and between light and dark areas. Fourth, a motion predictor was computed using a spatiotemporal Gabor filter model (Nishimoto et al., 2011). A detailed description of all predictor metrics can be found in the Methods section.

### Visual features robustly predict aesthetic appeal ratings

For each participant, we then used a cross-validation approach across movie segments to predict moment-to-moment aesthetic ratings from visual features. In all analyses, linear models were trained to predict aesthetic appeal ratings from the visual predictors on a subset of the movie segments and evaluated on a left-out movie segment. Specifically, for each cross-validation fold, one segment was held out as the test set, a second segment served as a validation set, and the remaining segments (seven for *Home*; six for *Loving Vincent*) were used for training. A ridge regression model was trained to predict aesthetic ratings (Y) from the four predictor groups (twelve visual predictors) (X), including visual fluency, symmetry, color statistics, and motion energy. Models were fitted on the training set with a range of regularization parameters λ, and predictive performance was evaluated on the validation set for each. We then selected the λ with the best performance on the validation set. Finally, a model with this optimal λ was fitted on the training set and in turn evaluated on the held-out test set. Model performance was quantified by computing the Pearson correlation between predicted and observed aesthetic appeal ratings. The model significantly predicted aesthetic appeal ratings in both *Home* (*t*(36) = 12.18, *p* < .001) and *Loving Vincent* (*t(*36) = 8.49, *p* < .001) (Fig. 3). These findings indicate that visual features predict dynamic aesthetic experiences and that predictions are robust for different visual content.

**Figure 3.**
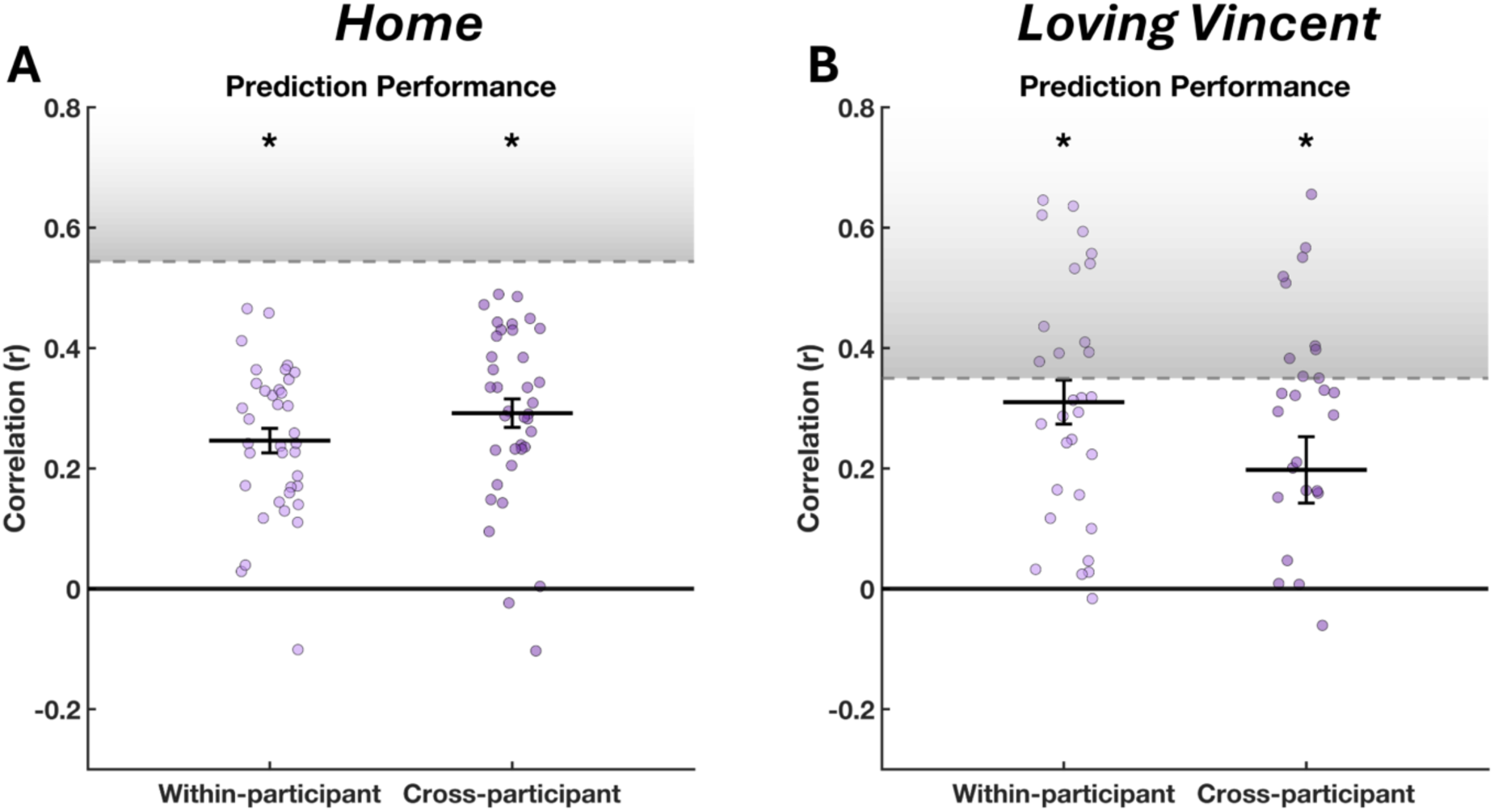
Within- and cross-subject prediction performance for *Home* (A) and *Loving Vincent* (B). The grey area indicates the noise ceiling, representing the maximum achievable prediction performance, computed by correlating each subject’s ratings with the average ratings of all other subjects across movie segments. Dots represent predictive performance in individual participants, quantified as the correlation between predicted and observed ratings. Error bars indicate SEM, asterisks indicate p < 0.05.

### Visual features predict aesthetic appeal across participants

The previous analysis, in which models are trained and tested within individual participants, demonstrates that visual features predict aesthetic appeal ratings within individuals. Next, we asked whether these relationships generalize across individuals, which would indicate that participants evaluate visual features in similar ways. To assess cross-subject generalization, we trained ridge regression models to predict group-level aesthetic dynamics from visual features and evaluated them on individual left-out participants. The regularization parameter (λ) was selected using leave-one-segment-out cross-validation on the group-mean data. In this cross-participant analysis, the model still showed significant predictive performance for both *Home* (*t(*36) = 12.35, *p* < .001) and *Loving Vincent* (*t(*29) = 3.58, *p* = .001) (Fig. 3). These findings demonstrate that visual feature models yield predictions that generalize across participants, suggesting that visual features contribute to aesthetic appeal in similar ways for different individuals. Yet cross-participant predictions were more accurate for *Home* than for *Loving Vincent*, consistent with greater individual variability in the aesthetic appeal of artistic stimuli (Vessel et al., 2018).

### Visual fluency and color statistics drive predictions of aesthetic experiences

After establishing that visual features predict aesthetic experience within and across participants, we next asked which features contribute most to this prediction. To address this, we conducted a model comparison analysis in which models were retrained while leaving out one predictor group (i.e., fluency, symmetry, color, motion) at a time. The change in performance (Δr) was computed by subtracting the reduced models’ predictive performance from the full model’s performance. We then performed one-sample right-tailed t-tests to determine whether excluding each predictor group significantly reduced model performance.

For *Home*, both within- and cross-participant analyses showed that leaving out visual fluency (within: *t*(36) = 4.59, *p* < .001; cross: *t*(36) = 4.17, *p* < .001), symmetry (within: *t*(36) = 2.76, *p* = .006; cross: *t*(36) = 7.24, *p* < .001), and color (within: *t*(36) = 8.43, *p* < .001; cross: *t*(36) = 12.62, *p* < .001) significantly reduced model performance, while leaving out motion had no effect (Fig. 4A). For *Loving Vincent,* leaving out visual fluency (within: *t*(29) = 4.32, *p* < .001; cross: *t*(29) = 4.16, *p* < .001) and color (within: *t*(29) = 4.45, *p* < .001; cross: *t*(29) = 1.16, trending at *p* = .10) reduced model performance, while leaving out symmetry and motion did not reduce model performance (Fig. 4B).

**Figure 4.**
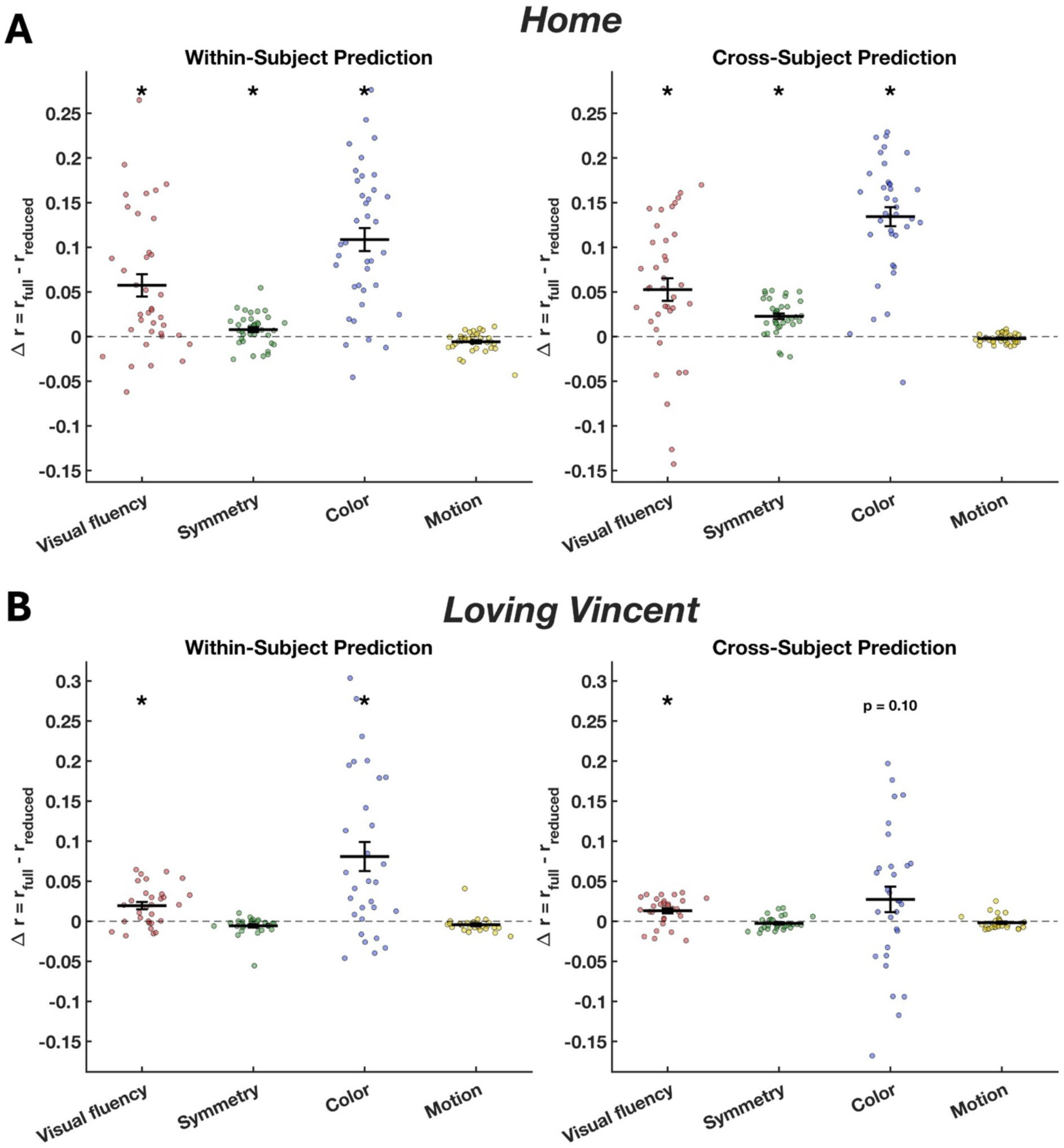
(A) Model comparison results for *Home*. The contribution of each predictor group to the performance of the full model is shown for both within- and cross-subject analyses. The y-axis represents the change in predictive performance (Δr), computed as the difference between the full and reduced models. Visual fluency, symmetry, and color predictors significantly reduced model performance in both within- and cross-participant analyses. (B) Model comparison results for *Loving Vincent.* Leaving out visual fluency and color predictors reduced model performance in both within- and cross-participant analyses. Error bars indicate SEM, asterisks indicate p < 0.05.

### Visual features predict aesthetic appeal across different movie content

Next, we tested whether models trained on one movie can predict aesthetic appeal ratings in the other movie, despite the marked differences in content and style across the two movies. This analysis provides a critical test of how visual features affect aesthetic appeal across domains. Here, models were trained on *N* − 1 segments of the average ratings and predictors from one movie (e.g., *Home*), using the remaining segment for regularization, and then retrained on all *N* segments. The final model was subsequently tested on each segment of each participant from the other movie (e.g., *Loving Vincent*).

Models trained on one movie significantly predicted aesthetic appeal ratings in the other movie, in both train-test directions (*Home* to *Loving Vincent*: *t*(29) = 2.9, *p* = .007; *Loving Vincent* to *Home*: *t*(36) = 14.75, *p* < .001) (Fig. 5). Predictive performance was less variable across individuals when the model was trained on *Loving Vincent*. This may reflect constraints in feature exploitation during training: models that use a narrower set of visual features when trained on *Loving Vincent* may yield lower prediction variability.

**Figure 5.**
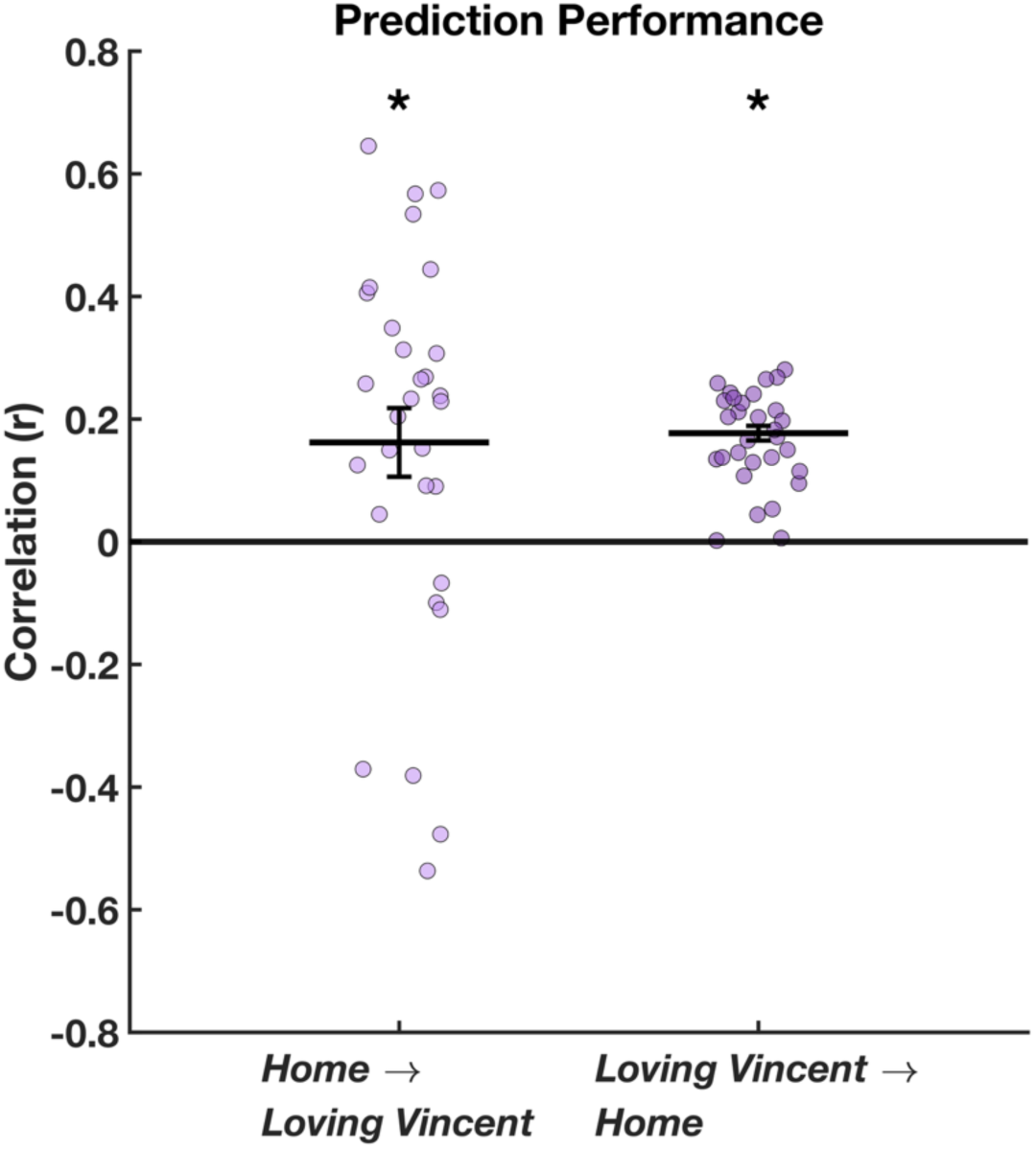
Cross-dataset prediction performance. Models were trained on the average aesthetic ratings of *Home* and tested on individual participants in *Loving Vincent*, and vice versa. Predictive performance is shown as the correlation between predicted and observed ratings. Error bars indicate SEM, asterisks indicate p < 0.05.

### Cross-movie predictions are predominantly driven by color features

To further examine the contribution of each predictor group to this cross-dataset generalization, we again performed a model comparison analysis. For each train-test direction, models were evaluated while leaving out one predictor group at a time. Prediction performance was again quantified as the correlation between predicted and observed ratings, and the change in performance (Δr) was computed by subtracting the reduced model performance from the full model performance. One-sample right-tailed *t*-tests were used to assess whether excluding each predictor group significantly reduced prediction performance.

When models were trained on the average aesthetic appeal ratings from *Home* and tested on the individual ratings in *Loving Vincent*, model comparison analysis showed that leaving out color (*t*(29) = 3.1, *p* = .005) and motion (*t*(29) = 3.07, *p* = .005) predictors significantly reduced model performance, whereas other predictor groups did not show significant effects (Fig. 6A). When the models were trained on the average ratings from *Loving Vincent* and tested on the individual ratings from *Home*, leaving out both color (*t*(36) = 12.8, *p* < .001) and motion (*t*(36) = 2.5, *p* = .02) predictors reduced model performance, whereas visual fluency and symmetry did not (Fig. 6B).

**Figure 6.**
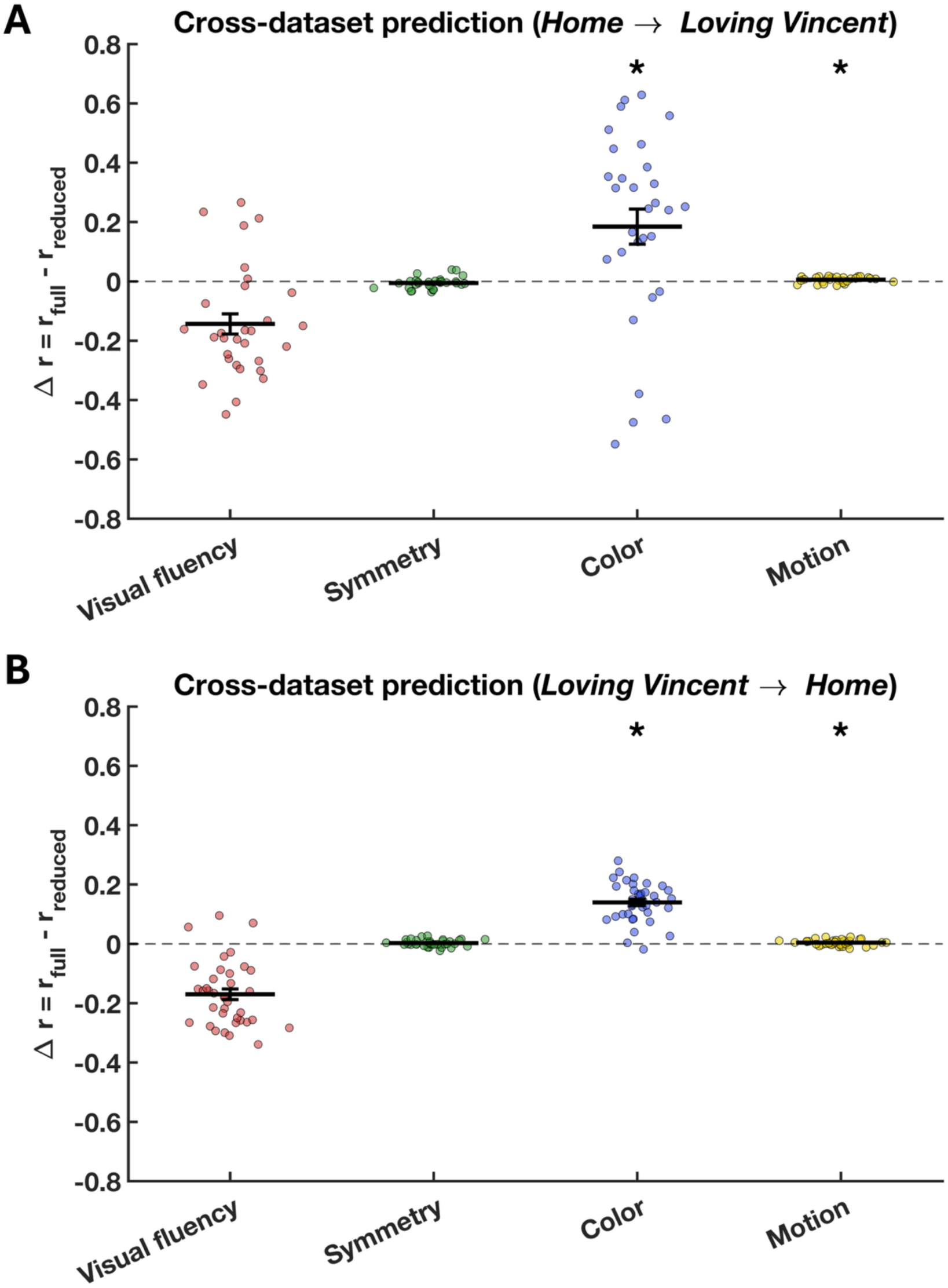
(A) Cross-dataset prediction performance (training on *Home*, testing on *Loving Vincent*) using model comparison. The y-axis shows the change in prediction performance (Δr), computed as the difference between the full and reduced model performances. Only color statistics significantly reduced model performance, whereas other predictor groups did not. (B) Cross-dataset prediction performance (training on *Loving Vincent*, testing on *Home*). Color statistics and motion energy significantly reduced model performance, whereas visual fluency and symmetry did not. Error bars indicate SEM, asterisks indicate p < 0.05.

Together, these results indicate that color statistics provide the most consistent contribution to cross-dataset prediction, suggesting that they capture shared constraints on aesthetic experience across visually and stylistically distinct movies. Excluding the motion predictor also yielded a significant reduction in performance. However, the effect was numerically very small and motion did not consistently contribute to predictions within individual movies. The effects observed here may thus stem from systematic but weak interactions with other predictors. Interestingly, while the visual fluency predictor group was informative for predictions within each movie, it did not generalize to the cross-dataset analysis. Specifically, in the naturalistic movie *Home*, similarity and integration yielded positive regression weights, whereas L2 norm yielded a negative weight, consistent with previous findings on natural scenes (Nara & Kaiser, 2024). In contrast, in the artistic movie *Loving Vincent*, L2 norm yielded positive weights, while similarity and integration yielded negative weights (Fig. S3). This may indicate differences in how visual fluency contributes to aesthetic experience across different types of visual content.

## Discussion

Our study demonstrates that aesthetic experience can be robustly predicted from visual features. By combining multiple low- and mid-level image-computable features, we modeled moment-to-moment fluctuations in perceived beauty across dynamic movie stimuli. These visual-feature models not only predicted aesthetic appeal ratings on an individual level; trained models also predicted aesthetic appeal across different observers and across the two movies. Our findings thus support the idea that aesthetic experience is rooted in perceptual processes, which play out in similar ways across individuals and different content.

### Visual features predict aesthetic appeal across observers

The finding that model predictions generalize across observers suggests that aesthetic judgments are not purely idiosyncratic but partly grounded in shared visual feature preferences. This finding challenges the common assumption that “beauty lies in the eye of the beholder,” instead pointing to a form of “perceptual” beauty that is shared across individuals. At the same time, predictions across observers varied with content: cross-participant predictive performance was reduced for *Loving Vincent* compared to *Home*, which suggests that the aesthetic appeal of artistic stimuli is more variable across individuals. This finding aligns with prior work showing that artworks and architecture tend to evoke more disagreement among people than natural scenes, likely due to differences in interpretation, exposure, and cultural artifacts (Vessel et al., 2018). Yet, reduced cross-participant predictions may not only relate to higher-level cognitive differences but also to individual variability in how visual features are mapped onto aesthetic appeal: Studies on the neural representation of aesthetic appeal indicate that even early brain responses associated with perceptual processing are partly idiosyncratic (Kaiser & Nyga, 2020; Kaiser, 2022).

### Color predicts aesthetic appeal across different content

Color statistics and visual fluency were the most informative predictors both within and across participants, whereas symmetry and visual motion played a comparatively minor role. Notably, color features were the only predictors that robustly generalized across movies. Among the color predictors we used, overall variation in colors (colorfulness), along with brightness contrast, contributed most to successful predictions of aesthetic ratings (Fig. S3), consistent with prior work showing that colorfulness is a significant predictor of aesthetic preference (Reinecke et al., 2013). The positive contribution of colorfulness (positive beta coefficient) indicates a preference for colorful visual inputs. Moreover, our results indicate that green and blue hues are associated with greater aesthetic appeal than red and yellow hues (Fig. S3). Consistent with this pattern, earlier work has shown that bluish and greenish color ranges tend to be preferred over darker or brownish tones in aesthetic evaluations of paintings (Amirshahi et al., 2016). Our results suggest that such color preferences may play out similarly across very different content (here: two different movies). This is in line with a previous study showing that preferences for color compositions persist even when spatial structure and semantic content are disrupted, such as in scrambled or patchwork images (Nakauchi et al., 2022). Interestingly, the same study reported only weak and non-replicable cultural differences, suggesting that the dominance of color statistics in aesthetic preference is robust across observers.

This notion is consistent with our cross-movie predictions being primarily driven by color features and indicates that color can serve as a reliable predictor of aesthetic appeal despite substantial higher-level differences in context, content, and cultural interpretation. The importance of color for aesthetic appeal is potentially rooted in biological or ecological factors, such as sensitivity to natural color distributions (Nascimento et al., 2021).

### Fluency predicts aesthetic appeal in nuanced ways

Although visual fluency was predictive within each movie, it showed qualitatively different contributions across the two movies. In the naturalistic movie *Home*, fluency-related measures aligned with previous findings on natural scenes: similarity and integration contributed positively to the predictions, indicating that efficient integration of image parts predisposes greater aesthetic appeal, whereas the L2 norm showed a negative contribution, indicating that representational sparseness predisposes greater aesthetic appeal (Nara & Kaiser, 2024; Tang et al., 2025). However, in *Loving Vincent*, the contribution of these three predictors was reversed (Fig. S3), potentially indicating that greater fluency yields *lower* aesthetic appeal. This divergence indicates that fluency effects may be modulated by the statistical structure and style of the visual input. One possible explanation is that more artistic depiction (like the painting style in *Loving Vincent*) engages a different mode of aesthetic evaluation, similar to the processing of artworks. In this mode, observers may shift from rapid, perceptually driven judgments to more reflective processing that enables them to understand or interpret more deeply and become more sensitive to complexity, ambiguity, and higher-level features (Graf & Landwehr, 2015). In this reflective mode, less fluent inputs may be more attractive, as they provide a richer basis for engagement and interpretation. Another explanation is that the deep neural network used to derive these features (VGG16 trained on Places365) is trained on natural images rather than artworks (Castellano & Vessio, 2021) and may therefore capture fluency in a way that is better aligned with natural scene statistics than with painterly representations. Regardless, it is important to note that fluency was a powerful predictor for aesthetic appeal in both movies. Yet, its role may depend on the context or the representational format of the stimulus.

### How “good” are models based on interpretable visual features?

Despite the success of our feature-based prediction models, a considerable portion of variance in aesthetic ratings remains unexplained. This likely reflects the influence of higher-level factors, such as semantic meaning, emotional engagement, cultural differences, and individual differences in personality or prior experience (Chatterjee & Vartanian, 2016). At the same time, our approach can be considered a starting point, where the inclusion of other, more complex feature predictors could further push the boundaries of predictive accuracy. In this context, it is important to note that our analysis deliberately focused on a set of relatively simple and interpretable predictors. It is possible that more comprehensive feature spaces could yield improved predictions (Ibarra et al., 2017). Indeed, recent work has shown that “brute-force” approaches, directly regressing deep neural network (DNN) activity patterns onto aesthetic ratings (Conwell et al., 2021; Nara & Kaiser, 2024), achieve high predictive performance. However, the models employed in such studies are essentially black boxes, with regressions on high-dimensional spaces of arbitrary DNN features offering limited insight into the specific features driving aesthetic experiences. Our approach is complementary to that, emphasizing interpretability to identify which visual properties meaningfully contribute to perceived beauty.

### Looking forward: continuous ratings of aesthetic appeal as an emerging method

The present study highlights the value of using dynamic, naturalistic stimuli to investigate aesthetic experiences, while allowing participants to provide continuous, moment-to-moment ratings of aesthetic appeal. Importantly, our findings demonstrate that principles established in studies using static images and individual ratings generalize to complex, dynamic stimuli assessed with continuous measures. This suggests that core perceptual drivers of aesthetic experience remain stable across stimulus formats and timescales. At the same time, our findings further showcase the viability of studying aesthetic experiences with continuous ratings, offering an ecologically valid and temporally fine-grained account of aesthetic experience (Ortega & Whitney, 2025). The continuous rating approach opens new opportunities for investigating the temporal dynamics of aesthetic experience (Isik & Vessel, 2019). For instance, it can be used to link fluctuations in aesthetic appeal to fluctuations in neural dynamics during movie watching or to test predictive models of how aesthetic appeal unfolds over time (Brielmann et al., 2024; Frascaroli et al., 2023).

## Conclusion

In sum, our findings suggest that aesthetic experience is best understood as shared subjectivity: while individual differences and contextual factors play an important role, they are rooted in shared perceptual mechanisms that can be captured through visual features. Future research could integrate low-level feature-based approaches with higher-level cognitive and affective factors to develop a more comprehensive model of aesthetic experience.

## Method

### Participants

The study comprised two experiments. In Experiment 1, a total of 37 participants with normal or corrected-to-normal vision participated (*M_age_* = 26.76, SD = 4.28; 21 females). In Experiment 2, 30 participants with normal or corrected-to-normal vision took part (*M_age_* = 26.28, *SD* = 6.11; 15 females). All participants provided written informed consent and were reimbursed at a rate of €10 per hour for their participation. The study was approved by the General Ethics Committee of Justus Liebig University Giessen (approval no. AZ25/22).

### Movie Stimulus

In Experiment 1, participants watched the documentary *Home* divided into nine segments (474.3, 534.8, 529.3, 535.2, 520.7, 537.6, 533.3, 523.3, and 357.7 seconds; mean duration = 505.1 seconds). *Home* includes a diverse range of visual content, from the dance of a whale to forest fires, capturing both pleasing and unpleasing scenes from all around the world. In Experiment 2, the animated movie *Loving Vincent* was shown in 8 segments (584.3, 527, 604.6, 620.1, 603.5, 565.1, 544.6, and 601.1 seconds; mean duration = 581.3 seconds). *Loving Vincent* depicts an animated interpretation of Vincent van Gogh’s paintings and life story. Both movies were presented without sound. The segments did not overlap, and each was cut at the end of a scene to preserve coherence within segments. In Experiment 1, participants watched the movie in one of 8 randomized sequences of segments, and in Experiment 2, participants watched the movie in 18 randomized sequences. The movies exhibit relatively weak narrative continuity, which is largely lost in the absence of sound. After collecting data from twelve participants in Experiment 1, we slightly improved movie clipping, removing some black screens, which lasted only around 2 minutes in total (3.38% of the total duration).

### Procedure

In both experiments, participants sat in a darkened room with their heads stabilized by a chin and forehead rest, positioned 68 cm from the screen. Stimuli were displayed on a monitor with a resolution of 1280 × 720 pixels and a refresh rate of 100 Hz. At this viewing distance, the stimuli subtended 28.7 × 16.4 degrees of visual angle. During the movie segments, participants rated aesthetic appeal using a scale presented below the movie, ranging from −144 to +144 in steps of 6 units (49 discrete levels). A practice session of continuous aesthetic rating was shown at the beginning of each experiment. The experiment was programmed using Psychtoolbox version 3.0.19 in MATLAB R2022a (MathWorks, Natick, MA, USA) on a Windows 10 PC.

### Behavioral data processing

For each participant, we computed the average reaction-time delay in the continuous rating task and adjusted their rating time series accordingly. To accomplish this, movie cuts were identified by calculating frame-to-frame pixel differences. We then measured the average time to the first rating change following each cut and used this latency to temporally shift the rating data, with the shift limited at a maximum of two seconds.

Furthermore, the first two seconds of each movie segment were excluded, as participants required additional time to adjust their ratings at segment onsets.

### Image-computable predictors

The following image-computable visual features were extracted for each movie frame. The values were then averaged across 25-frame (i.e., 1s) chunks. This was done because faster changes would probably not be reflected prominently in an adjustment of the rating slider.

### DNN-based measures of visual fluency

This set of predictors features three measures: Part-similarity, integration, and L2 norm, all derived from DNN representations (Nara & Kaiser, 2024). We employed a VGG16 deep convolutional neural network (Simonyan & Zisserman, 2015) trained for scene classification on the Places365 dataset (Zhou et al., 2018). The pretrained network weights were obtained from the official Places365 repository and were originally implemented in Caffe (Jia et al., 2014). The model was subsequently imported into MATLAB using the Deep Learning Toolbox Caffe importer.

We presented the network with either the full movie frame or spatially partitioned versions of the frame. The partitions were generated by dividing each movie frame into an 8 × 8 grid of equal-sized squares. Two complementary image sets were then created: one containing all odd-numbered squares and the other containing all even-numbered squares, analogous to the black and white fields of a checkerboard pattern. The full image and both partitioned versions were fed separately into the network, and layer-specific activation patterns were extracted from the 11th layer, as previous work indicated that this layer yields the most robust predictions of aesthetic appeal (Nara & Kaiser, 2024).

Part-similarity was quantified by correlating the activation patterns of the two complementary halves. Integration was assessed by averaging the activation vectors of the two halves and correlating this mean activation with the activation elicited by the full image. The resulting correlation values were sign-inverted so that weaker similarity between parts and whole reflected higher integration. Finally, L2 norms were computed by summing the squared activations of all units in the selected layer to quantify overall activation strength. Fig. S4 shows a conceptual illustration of the three DNN-based measures.

### Symmetry

Symmetry was quantified by calculating pixel-wise correlations across all possible horizontal and vertical axes of reflection. For each image, the maximum correlation value, weighted by the number of pixels contributing to the computation, was taken as the measure of horizontal and vertical symmetry (Damiano et al., 2023).

### Color statistics

Colorfulness, greenness, and blueness were computed in CIELab color space. For each frame, we extracted the *a* (green-red) and *b* (blue-yellow) channels. The mean and standard deviation of the *a* and *b* channels were computed across all pixels to quantify chromatic bias and distribution. The colorfulness was defined as the Euclidean norm of the *a* and *b* standard deviations plus 0.3 times the Euclidean norm of their mean values, capturing both chromatic variability and overall color intensity (Hasler & Suesstrunk, 2003). Greenness and blueness were defined as sign-inverted versions of the averages of the *a* and *b* channels, respectively.

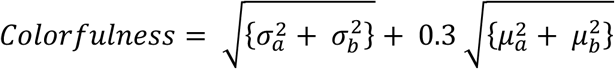

Mean saturation was computed as the average of the HSL color space saturation values across all pixels for each frame. Hue contrast measures how far apart the main colors in an image are, ignoring very dull or overly bright/dark pixels; higher values mean the colors are more spread out. Brightness contrast was computed in RGB color space as the width of the smallest symmetric histogram interval centered on the most frequent brightness bin that contained 98% of all pixels (Iigaya et al., 2021).

### Motion Energy

Motion energy was quantified using a spatiotemporal Gabor filter model, implemented with the publicly available MATLAB code from https://github.com/gallantlab/motion_energy_matlab (Nishimoto et al., 2011). The motion data was transformed using Principal Component Analysis (PCA) to capture its main patterns of variation. Only the first principal component is retained, representing the dominant motion signal, which is then stored as a 1D motion energy predictor.

### Ridge regression models

To predict continuous aesthetic ratings from visual features, we applied ridge regression separately for each participant. We implemented a fully exhaustive nested cross-validation procedure across all movie segments (nine segments for *Home*, and eight segments for *Loving Vincent*). For each fold, one segment served as the test set, one segment as the validation set for hyperparameter selection, and the remaining segments as the training set. This resulted in 72 folds per participant (9 test segments × 8 validation segments) for *Home*, and 56 folds for *Loving Vincent* (8 test segments × 7 validation segments). Within each fold, predictor variables were standardized based on the training data. Ridge models were fitted on the training set for each λ value, and the optimal λ was chosen from a logarithmically spaced range (10⁻⁵ to 10⁵, 100 values) by minimizing mean squared error on the validation set. The final model with the selected λ was then evaluated on the held-out test set.

Model performance was quantified using Pearson correlation between predicted ratings and observed ratings for each segment. Performance was averaged across all folds per participant for both movies. At the group level, mean performance across participants was tested against zero using one-sample t-tests. To examine predictor contributions, regression coefficients were averaged across folds within each participant. Group-level significance of coefficients was assessed using one-sample t-tests across participants, with false discovery rate (FDR) correction applied to account for multiple comparisons.

### Cross-participant predictions

To assess cross-subject generalization, we trained ridge regression models to predict leave-one-subject-out group-mean rating time courses from the selected predictors and then evaluated these models against the ratings of the left-out participant. For each left-out participant and movie segment, the training target was the mean rating time course of the remaining *N* − 1 participants. The regularization parameter (λ) was selected separately for each participant and model using leave-one-segment-out cross-validation: in each fold, the model was trained on all but one segment and validated on the held-out segment by minimizing mean squared error. Using the optimal λ, the final model was fit across all segments and then used to predict the left-out participant’s moment-to-moment ratings for each segment. Prediction accuracy was quantified as the Pearson correlation between predicted and observed ratings, and correlations were averaged across segments to obtain a single performance estimate per participant. Group-level significance was assessed using one-sample t-tests across participants, with false discovery rate (FDR) correction.

### Assessing the relative contribution of features

The same ridge regression procedure was repeated with reduced models, excluding one predictor group each time. Then, the correlations of predicted and observed ratings for each reduced model were subtracted from the correlations obtained with the full model. We conducted a one-tailed t-test with FDR correction to compare whether the predictor group significantly reduced the model performance and contributed to predicting aesthetic preferences for both movies.

### Cross-movie predictions

Next, we examined whether models trained on aesthetic ratings for one type of content could predict ratings for a different type of content. To this end, we constructed two ridge regression models. In each case, models were trained on N−1 segments of the average ratings and predictors from one movie (e.g., *Home*), using the remaining segment for regularization, and then retrained on all N segments. The final model was subsequently tested on each segment of each participant from the other movie (e.g., *Loving Vincent*). The same procedure was applied in the reverse direction (training on *Loving Vincent* and testing on *Home*). Further, we again performed model comparison analyses to examine how cross-dataset predictions are driven by the different predictor groups.

## Acknowledgements

Author contributions

M.A.E and D.K. designed research; M.A.E and N.B. performed research; M.A.E analyzed data; D.K. provided equipment and advice on study design and analysis; M.A.E and D.K. wrote the paper.

## Funding acquisition

This work was supported by the Deutsche Forschungsgemeinschaft (DFG), KA4683/6-1 (project no. 536053998); and under Germany’s Excellence Strategy (EXC 3066/1, “The Adaptive Mind”, project no. 533717223). It was further supported by a European Research Council (ERC) Starting Grant (PEP, ERC-2022-STG 101076057). Views and opinions expressed are those of the authors only and do not necessarily reflect those of the funders. Neither the funders nor the granting authority can be held responsible for them.

## Competing interests

The authors declare no competing interests.

## Data availability

Data and code are available through the project’s OSF page (https://osf.io/e2qk8).

## Supporting Information

### Correlation

**Figure S1.**
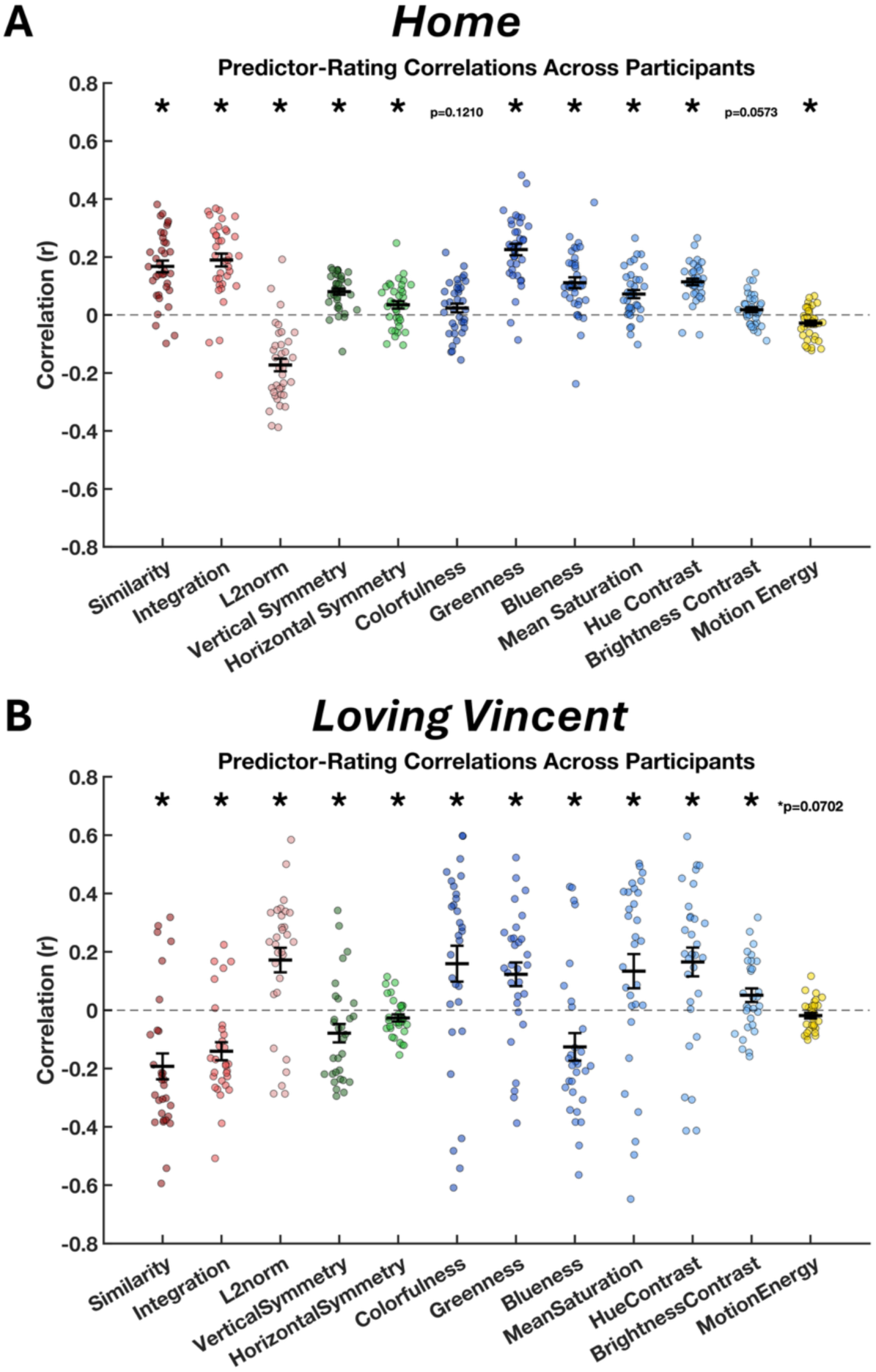
Correlations between individual predictors and aesthetic ratings. In addition to the regression models, we computed correlations between each predictor and the continuous aesthetic ratings. Group-level one-sample t-tests were performed with FDR correction to assess the relationship between each predictor and dynamic aesthetic experience. Error bars indicate SEM, asterisks indicate p < 0.05.

### Partial correlation

**Figure S2.**
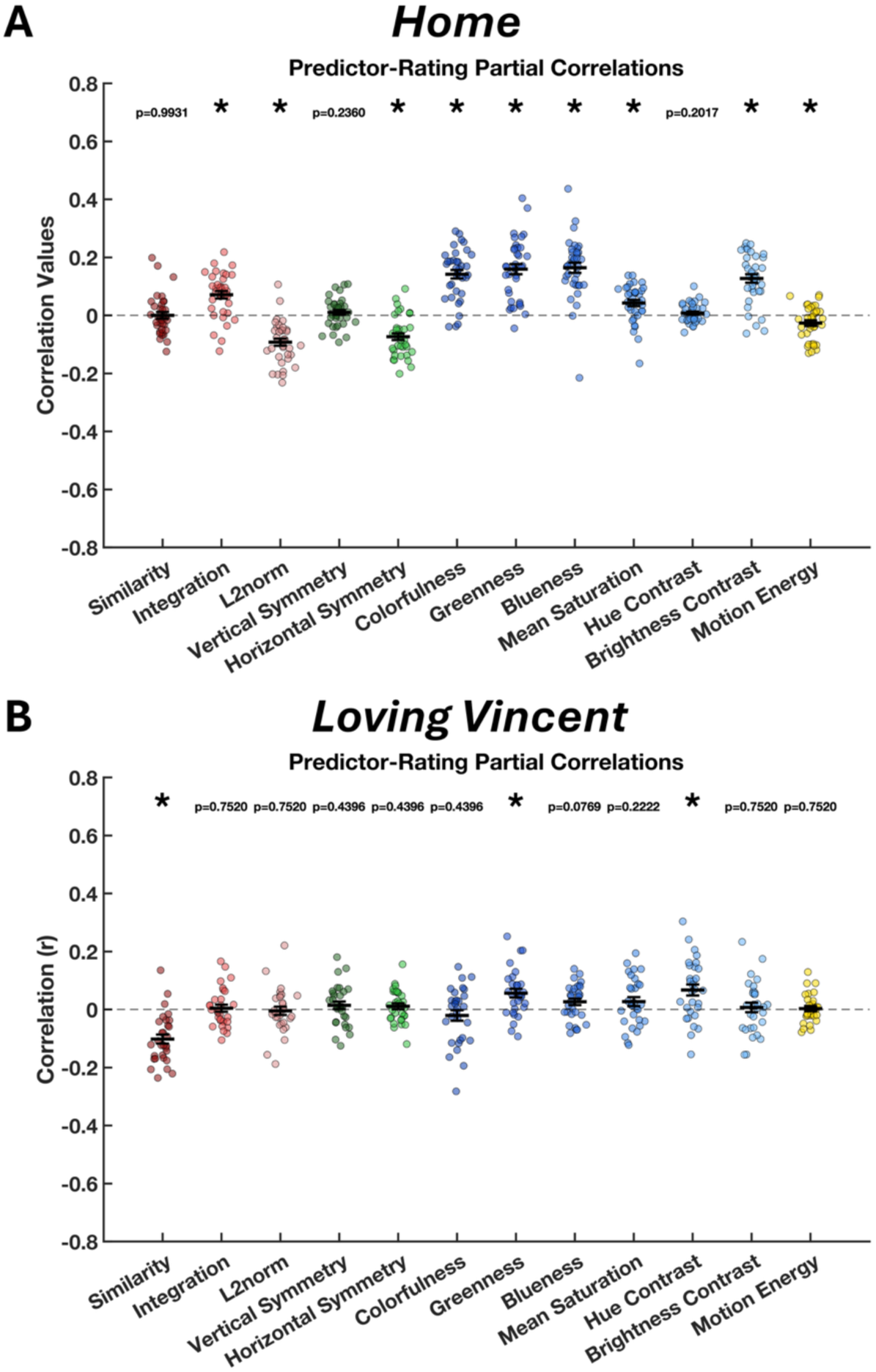
Partial correlations between individual predictors and aesthetic ratings. Partial correlations were computed to examine the relationship between each predictor and aesthetic ratings while controlling for the influence of all other predictors. Error bars indicate SEM, asterisks indicate p < 0.05.

### Beta coefficients

**Figure S3.**
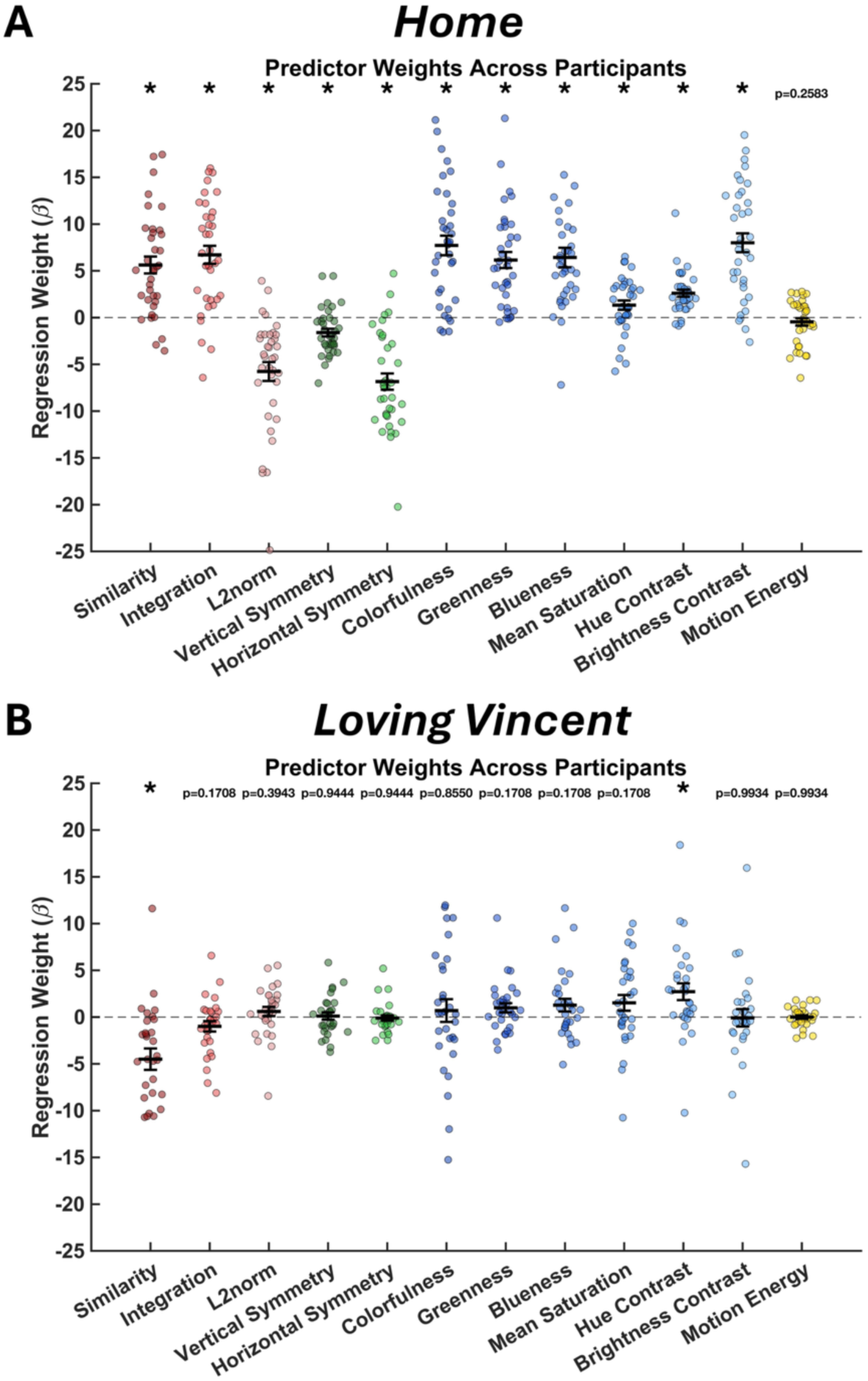
Regression coefficients (beta weights) of predictors from the regression models. These values reflect the unique contribution of each predictor to explaining variance in aesthetic ratings. Group-level one-sample t-tests were performed on the beta coefficients, with FDR correction applied for multiple comparisons. Error bars indicate SEM, asterisks indicate p < 0.05.

**Figure S4.**
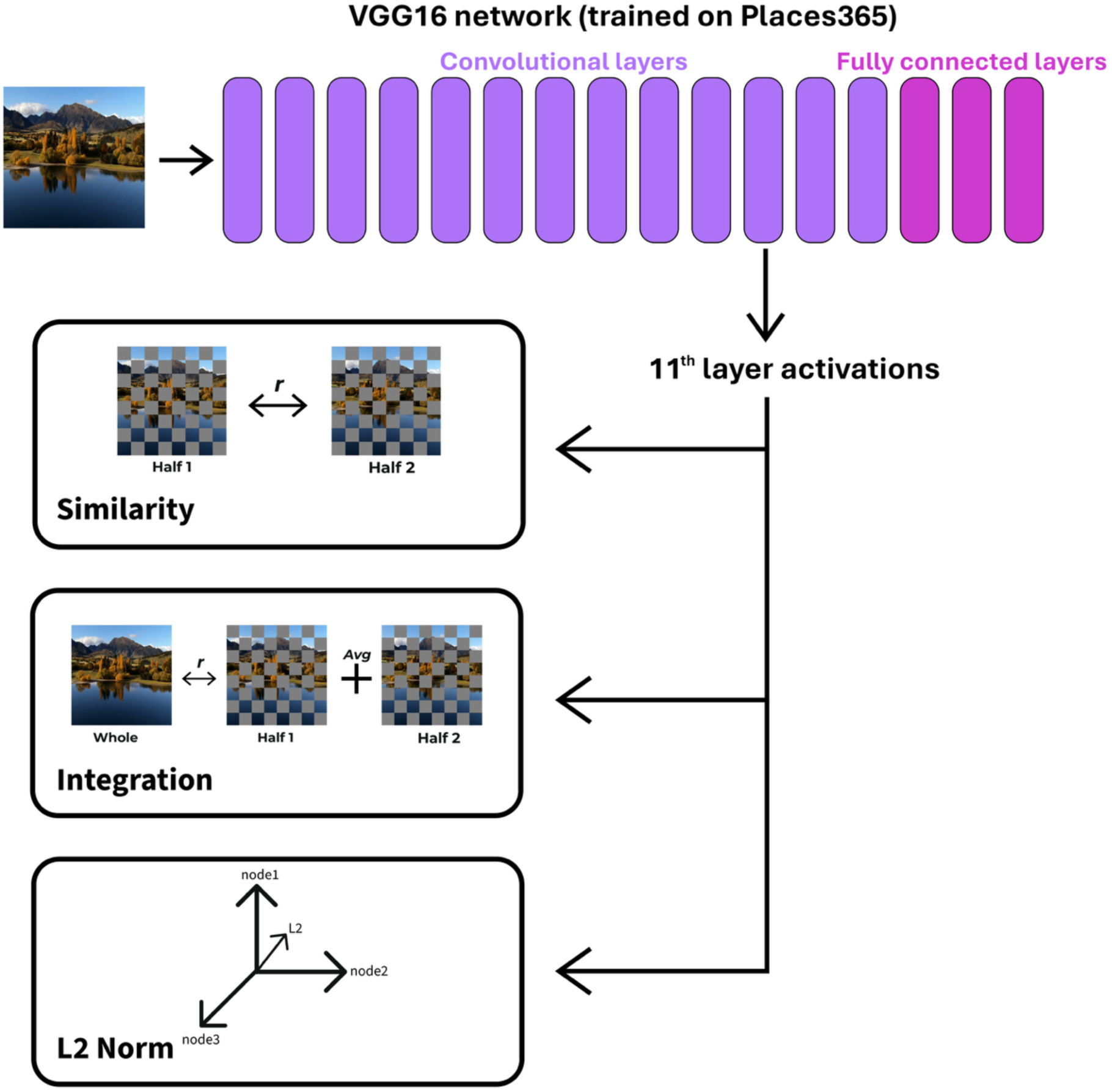
Coding efficiency predictors derived from a deep neural network (VGG16). For each frame, full and two checkerboard-split (8 x 8) image versions were processed, and activations from the 11th layer were extracted. Similarity was computed as the correlation between activations of the two halves, and integration as the correlation between the full image and the average of the two halves (the sign was inverted so higher values reflect greater integration). The L2 norm was computed from the activation vectors.

## Notes

https://osf.io/e2qk8

### Competing Interest Statement

The authors have declared no competing interest.

## References

Amirshahi, S. A., Hayn-Leichsenring, G. U., Denzler, J., & Redies, C. (2016). Color: A crucial factor for aesthetic quality assessment in a subjective dataset of paintings. arXiv preprint arXiv:1609.05583.

Brielmann, A. A., & Dayan, P. (2022). A computational model of aesthetic value. Psychological Review, 129(6), 1319.

Brielmann, A. A., Berentelg, M., & Dayan, P. (2024). Modelling individual aesthetic judgements over time. Philosophical Transactions of the Royal Society B: Biological Sciences, 379(1895).

Carbon, C. C. (2010). The cycle of preference: Long-term dynamics of aesthetic appreciation. Acta psychologica, 134(2), 233–244.

Castellano, G., & Vessio, G. (2021). Deep learning approaches to pattern extraction and recognition in paintings and drawings: an overview. Neural Computing and Applications, 33(19), 12263–12282.

Chatterjee, A. (2011). Neuroaesthetics: A Coming of Age Story. Journal of Cognitive Neuroscience, 23(1), 53–62. 10.1162/jocn.2010.21457

Chatterjee, A., & Vartanian, O. (2016). Neuroscience of aesthetics. Annals of the New York Academy of Sciences, 1369(1), 172–194.

Conwell, C., Graham, D., & Vessel, E. A. (2021). The Perceptual Primacy of Feeling: Affectless machine vision models robustly predict human visual arousal, valence, and aesthetics.

Corradi, G., Belman, M., Currò, T., Chuquichambi, E. G., Rey, C., & Nadal, M. (2019). Aesthetic sensitivity to curvature in real objects and abstract designs. Acta Psychologica, 197, 124–130.

Damiano, C., Wilder, J., Zhou, E. Y., Walther, D. B., & Wagemans, J. (2023). The role of local and global symmetry in pleasure, interest, and complexity judgments of natural scenes. Psychology of Aesthetics, Creativity, and the Arts, 17(3), 322–337. 10.1037/aca0000398

Dhar, S., Ordonez, V., & Berg, T. L. (2011, June). High level describable attributes for predicting aesthetics and interestingness. In CVPR 2011 (pp. 1657-1664). IEEE.

Enquist, M., & Arak, A. (1994). Symmetry, beauty and evolution. Nature, 372(6502), 169– 172. 10.1038/372169a0

Farzanfar, D., & Walther, D. B. (2023). Changing What You Like: Modifying Contour Properties Shifts Aesthetic Valuations of Scenes. Psychological Science, 34(10), 1101–1120. 10.1177/09567976231190546

Fechner, G. T. (1876). Vorschule der Aesthetik. Breitkopf & Härtel.

Frascaroli, J., Leder, H., Brattico, E., & Van de Cruys, S. (2024). Aesthetics and predictive processing: grounds and prospects of a fruitful encounter. Philosophical Transactions of the Royal Society B, 379(1895), 202

Graf, L. K., & Landwehr, J. R. (2015). A dual-process perspective on fluency-based aesthetics: The pleasure-interest model of aesthetic liking. Personality and social psychology review, 19(4), 395–410.

Graham, D., Schwarz, B., Chatterjee, A., & Leder, H. (2016). Preference for luminance histogram regularities in natural scenes. Vision Research, Vision and the Statistics of the Natural Environment, 120, 11–21. 10.1016/j.visres.2015.03.018

Hasler, D., & Suesstrunk, S. E. (2003). Measuring colorfulness in natural images. Human Vision and Electronic Imaging VIII, 5007, 87–95. 10.1117/12.477378

Hekkert, P., Snelders, D., & Van Wieringen, P. C. (2003). ‘Most advanced, yet acceptable’: Typicality and novelty as joint predictors of aesthetic preference in industrial design. British journal of Psychology, 94(1), 111–124.

Huang, Y., Xue, X., Spelke, E., Huang, L., Zheng, W., & Peng, K. (2018). The aesthetic preference for symmetry dissociates from early-emerging attention to symmetry. Scientific Reports, 8(1), 6263. 10.1038/s41598-018-24558-x

Ibarra, F. F., Kardan, O., Hunter, M. R., Kotabe, H. P., Meyer, F. A. C., & Berman, M. G. (2017). Image Feature Types and Their Predictions of Aesthetic Preference and Naturalness. Frontiers in Psychology, 8. 10.3389/fpsyg.2017.00632

Iigaya, K., Yi, S., Wahle, I. A., Tanwisuth, K., & O’Doherty, J. P. (2021). Aesthetic preference for art can be predicted from a mixture of low- and high-level visual features. Nature Human Behaviour, 5(6), 743–755. 10.1038/s41562-021-01124-6

Isik, A. I., & Vessel, E. A. (2019). Continuous ratings of movie watching reveal idiosyncratic dynamics of aesthetic enjoyment. PLoS One, 14(10), e0223896.

Jia, Y., Shelhamer, E., Donahue, J., Karayev, S., Long, J., Girshick, R., Guadarrama, S., & Darrell, T. (2014). Caffe: Convolutional Architecture for Fast Feature Embedding. Proceedings of the 22nd ACM International Conference on Multimedia, MM ’14, 675–678. 10.1145/2647868.2654889

Jones, B. C., DeBruine, L. M., & Little, A. C. (2007). The role of symmetry in attraction to average faces. Perception & Psychophysics, 69(8), 1273–1277. 10.3758/BF03192944

Kaiser, D., & Nyga, K. (2020). Tracking cortical representations of facial attractiveness using time-resolved representational similarity analysis. Scientific reports, 10(1), 16852.

Kaiser, D. (2022). Characterizing dynamic neural representations of scene attractiveness. Journal of Cognitive Neuroscience, 34(10), 1988–1997.

Kirk, U., Skov, M., Hulme, O., Christensen, M. S., & Zeki, S. (2009). Modulation of aesthetic value by semantic context: An fMRI study. NeuroImage, 44(3), 1125–1132. 10.1016/j.neuroimage.2008.10.009

Leder, H., Belke, B., Oeberst, A., & Augustin, D. (2004). A model of aesthetic appreciation and aesthetic judgments. British Journal of Psychology, 95(4), 489–508. 10.1348/0007126042369811

Leder, H., Carbon, C.-C., & Ripsas, A.-L. (2006). Entitling art: Influence of title information on understanding and appreciation of paintings. Acta Psychologica, 121(2), 176–198. 10.1016/j.actpsy.2005.08.005

Leder, H., & Nadal, M. (2014). Ten years of a model of aesthetic appreciation and aesthetic judgments: The aesthetic episode – Developments and challenges in empirical aesthetics. British Journal of Psychology, 105(4), 443–464. 10.1111/bjop.12084

Lin, Y., Wagemans, J., & Op de Beeck, H. (2025). Using artificial neural networks to understand fluency in the perception of paintings. Proceedings of the Conference on Cognitive Computational Neuroscience (CCN).

Mather, K. B., Aleem, H., Rhee, Y., & Grzywacz, N. M. (2023). Social groups and polarization of aesthetic values from symmetry and complexity. Scientific Reports, 13(1), 21507. 10.1038/s41598-023-47835-w

Nakauchi, S., Kondo, T., Kinzuka, Y., Taniyama, Y., Tamura, H., Higashi, H., … & Nascimento, S. M. (2022). Universality and superiority in preference for chromatic composition of art paintings. Scientific Reports, 12(1), 4294.

Nascimento, S. M., Albers, A. M., & Gegenfurtner, K. R. (2021). Naturalness and aesthetics of colors–Preference for color compositions perceived as natural. Vision Research, 185, 98–110.

Nara, S., & Kaiser, D. (2024). Integrative processing in artificial and biological vision predicts the perceived beauty of natural images. ScieNce AdvANceS.

Nishimoto, S., Vu, A. T., Naselaris, T., Benjamini, Y., Yu, B., & Gallant, J. L. (2011). Reconstructing Visual Experiences from Brain Activity Evoked by Natural Movies. Current Biology, 21(19), 1641–1646. 10.1016/j.cub.2011.08.031

Ortega, J., & Whitney, D. (2025). Continuous affect tracking reveals that overestimation during the recollection of affect is idiosyncratic and stable. Journal of Vision, 25(11), 14–14.

Palmer, S. E., Schloss, K. B., & Sammartino, J. (2013). Visual Aesthetics and Human Preference. Annual Review of Psychology, 64(Volume 64, 2013), 77–107. 10.1146/annurev-psych-120710-100504

Perrett, D. I., Burt, D. M., Penton-Voak, I. S., Lee, K. J., Rowland, D. A., & Edwards, R. (1999). Symmetry and Human Facial Attractiveness. Evolution and Human Behavior, 20(5), 295–307. 10.1016/S1090-5138(99)00014-8

Reber, R., Schwarz, N., & Winkielman, P. (2004). Processing fluency and aesthetic pleasure: Is beauty in the perceiver’s processing experience?. Personality and social psychology review, 8(4), 364–382.

Redies, C. (2015). Combining universal beauty and cultural context in a unifying model of visual aesthetic experience. Frontiers in Human Neuroscience, 9. 10.3389/fnhum.2015.00218

Reinecke, K., Yeh, T., Miratrix, L., Mardiko, R., Zhao, Y., Liu, J., & Gajos, K. Z. (2013, April). Predicting users’ first impressions of website aesthetics with a quantification of perceived visual complexity and colorfulness. In Proceedings of the SIGCHI conference on human factors in computing systems (pp. 2049–2058).

Rhodes, G., Proffitt, F., Grady, J. M., & Sumich, A. (1998). Facial symmetry and the perception of beauty. Psychonomic bulletin & review, 5(4), 659–669.

Schloss, K. B., & Palmer, S. E. (2011). Aesthetic response to color combinations: Preference, harmony, and similarity. Attention, Perception, & Psychophysics, 73(2), 551–571. 10.3758/s13414-010-0027-0

Silvia, P. J., & Barona, C. M. (2009). Do People Prefer Curved Objects? Angularity, Expertise, and Aesthetic Preference. Empirical Studies of the Arts, 27(1), 25–42. 10.2190/EM.27.1.b

Simonyan, K., & Zisserman, A. (2015). Very Deep Convolutional Networks for Large-Scale Image Recognition (arXiv:1409.1556). arXiv. http://arxiv.org/abs/1409.1556

Swartz, A., Skelton, A. E., Mather, G., Bosten, J. M., Maule, J., & Franklin, A. (2024). The perceived beauty of art is not strongly calibrated to the statistical regularities of real-world scenes. Scientific Reports, 14(1), 19368. 10.1038/s41598-024-69689-6

Tang, Y., Cunningham, W. A., & Walther, D. B. (2025). Less is more: Aesthetic liking is inversely related to metabolic expense by the visual system. PNAS nexus, 4(12), pgaf347.

Oppenheimer, D. M. (2008). The secret life of fluency. Trends in cognitive sciences, 12(6), 237–241.

Van Geert, E., & Wagemans, J. (2020). Order, complexity, and aesthetic appreciation. Psychology of Aesthetics, Creativity, and the Arts, 14(2), 135–154. 10.1037/aca0000224

Van Geert, E., & Wagemans, J. (2024). Prägnanz in visual perception. Psychonomic Bulletin & Review, 31(2), 541–567.

Vartanian, O., Farzanfar, D., Munar, E., Skov, M., Hayn-Leichsenring, G., Ho, P. K., & Walther, D. B. (2024). Neural dissociation between computational and perceived measures of curvature. Scientific Reports, 14(1), 26529. 10.1038/s41598-024-76931-8

Vessel, E. A., & Rubin, N. (2010). Beauty and the beholder: Highly individual taste for abstract, but not real-world images. Journal of Vision, 10(2), 18. 10.1167/10.2.18

Vessel, E. A., Starr, G. G., & Rubin, N. (2012). The brain on art: Intense aesthetic experience activates the default mode network. Frontiers in Human Neuroscience, 6. 10.3389/fnhum.2012.00066

Vessel, E. A., Maurer, N., Denker, A. H., & Starr, G. G. (2018). Stronger shared taste for natural aesthetic domains than for artifacts of human culture. Cognition, 179, 121–131.

Zhou, B., Lapedriza, A., Khosla, A., Oliva, A., & Torralba, A. (2018). Places: A 10 Million Image Database for Scene Recognition. IEEE Transactions on Pattern Analysis and Machine Intelligence, 40(6), 1452–1464. 10.1109/TPAMI.2017.2723009

